# Pre-implantation mouse embryo movement under hormonally altered conditions

**DOI:** 10.1101/2021.11.19.469319

**Authors:** Hannah Lufkin, Diana Flores, Zachary Raider, Manoj Madhavan, Madeline Dawson, Anna Coronel, Dhruv Sharma, Ripla Arora

**Affiliations:** Department of Obstetrics Gynecology and Reproductive Biology, Michigan State University; Institute for Quantitative Health Science and Engineering, Michigan State University; Center for Statistical Training & Consulting, Michigan State University

**Author notes:** These authors contributed equally (Hannah Lufkin, Diana Flores). To whom correspondence should be addressed: **Ripla Arora**, Institute for Quantitative Health Science and Engineering, Department of Obstetrics, Gynecology and Reproductive Biology, 775 Woodlot Drive, Rm#3312, East Lansing, MI-48824, **Email:**.

## Abstract

Precise regulation of embryo movement is crucial to successful implantation, but the role of ovarian hormones in this process is not understood. We ascertain the effects of altered hormonal environment on embryo movement using two delayed implantation models: Natural lactational Diapause (ND), a naturally occurring alternate model of pregnancy, and Artificially induced Diapause (AD), a laboratory version of ND. Our previous work suggests that embryos in a natural pregnancy (NP) first display *unidirectional clustered embryo movement*, followed by *bidirectional scattering and spacing movement*. In contrast, in the ND model, embryos are present as clusters near the oviductal-uterine junction for ~24-hours longer than NP, followed by locations consistent with a *unidirectional scattering and spacing movement*. Intriguingly, the AD model closely resembles embryo location in NP and not ND. Further, unlike the popular paradigm of reduced estrogen (E2) levels in diapause E2 levels are comparable across NP, ND, and AD, while progesterone (P4) levels are reduced in ND and highly increased in AD when compared to NP. Exogenous administration of E2 or P4 modifies the unidirectional clustered embryo movement, while E2 treatment causes a reduction in P4 and affects the bidirectional phase of embryo movement. Taken together, our data suggest embryo movement can be modulated by both P4 and E2. Understanding natural hormonal adaptation in diapause provides an opportunity to determine key players regulating embryo movement and implantation success. This knowledge can be leveraged to understand pregnancy survival and implantation success in hormonally altered conditions in the clinic.

## INTRODUCTION

Ovarian hormones estrogen (E2) and progesterone (P4) prepare a receptive uterus for embryo attachment (Psychoyos, 1973). E2 levels are high during ovulation in both mice and humans, followed by increasing levels of P4 derived from the corpus luteum should a pregnancy ensue. A nidatory surge of E2 concomitant with high P4 induces implantation in mice (McCormack and Greenwald, 1974; Wang and Dey, 2006), although ovarian E2 is not needed for implantation in other species such as pigs, guinea pigs, rabbits, and hamsters (Cha and Dey, 2014). Whole-body deletion or uterine epithelial-specific deletion of both E2 receptors and P4 receptors disrupt the state of receptivity (Dupont et al., 2000; Franco et al., 2012; Lydon et al., 1995; Winuthayanon et al., 2010), supporting the essential role of hormones in this process. While hormonal control of receptivity has been studied in-depth, details on hormonal regulation of embryo transport in the uterus are sparse. Hormonal regulation of egg transport suggests an influence of both E2 and P4 in regulating contractions and tubal (oviductal) transport (Bylander et al., 2015). In the mouse oviduct, treatment with a P4 antagonist halts tubal egg movement, although the presence of P4 rather than the absolute levels of P4 seem to be important (Kendle and Lee, 1980). Prostaglandin, PGF2α treatment disrupts the corpus lutea and thus the luteal P4 and abrogates embryo mobility in the mare (Kastelic et al., 1987). In the rabbit, P4 plays a role in embryo movement by toning the muscle contractions (Boving, 1956). Uterine embryo movement in response to E2 has not been investigated extensively, likely because levels of E2 are high during ovulation but basal or low during embryo movement (Ma et al., 2003). An assessment of hormonal regulation of embryo movement is necessary to provide insights into how altered hormonal profiles in clinical scenarios such as hyperstimulation in artificial reproductive technologies (Gidley-Baird et al., 1986) or increased E2 in women with polycystic ovarian syndrome (van Houten and Visser, 2014) result in less optimal implantation with an unsuccessful pregnancy.

Naturally existing alterations in hormonal conditions represent evolutionarily selected regimens that support pregnancy success despite variation in hormone levels. One such example of this phenomenon in mammals is diapause. Diapause is a delay in implantation, induced in response to stress that leads to modification of the embryo growth and the uterine environment to pause pregnancy and prevent implantation until conditions are more optimal for survival of the young at birth (Cha and Dey, 2014; Fenelon et al., 2014; Mead, 1993). Two types of diapause occur naturally in the wild: obligate and facultative diapause. In obligate diapause, which occurs in the mink, skunk, bears, and some wallabies, every pregnancy is paused to align the birth of the offspring with a favorable season for survival (Lopes et al., 2004). In both mink and the skunk, compared to post-implantation gestation, P4 levels are reduced during diapause (Douglas et al., 1998; Mead, 1981). Facultative diapause is induced by external factors such as metabolic stress, access to resources, or lactation and occurs in rodents (rats and mice) and marsupials (wallabies and kangaroos) (Lopes et al., 2004). In lactational diapause, termed here natural diapause (ND), lactation by the first litter regulates ovarian hormone levels to delay embryo attachment. Mechanistically, in the Tammar wallaby, suckling young cause prolactin secretion, which suppresses the activity of the corpus luteum causing low levels of P4. Prolactin levels decrease when the suckling pouch with the pups is removed, so the corpus luteum and blastocyst activity resume (Renfree and Shaw, 2014). Female mice enter post-partum estrus upon delivering the first litter, and this rise in E2 leads to a mating event causing lactational diapause. Unlike the mink, skunk, and wallaby, lactating mice are thought to have an active corpus luteum (Whitten, 1958), and it is proposed that continued P4 production from the corpus luteum (in the absence of E2) maintains the pregnancy in a paused state while preventing embryo implantation (Mead, 1993). Further, studies where a single injection of E2 can induce implantation in lactating mice has led to a supposition that P4 levels are normal and E2 levels are lower in ND model of pregnancy (McLaren, 1968). Ovarian weights of lactating mice are lower than nonlactating mice (Whitten, 1955), indicating that although the corpus lutea are active, P4 production could be affected in mice undergoing lactational diapause. In support of this idea, exogenous P4 injection in lactating mice or rats, similar to E2, can induce implantation in rats and mice (McLaren, 1971; Yoshinaga, 1961), but levels of E2 and P4 have not been systematically assessed in the ND model of mouse pregnancy.

Diapause is an important model for understanding how the uterine environment and the embryo are modulated to delay pregnancy naturally in the mouse, but generating this model in the laboratory is time-consuming and costly. Thus, an induced delay model is more commonly used to study diapause in the lab setting. In this model termed here artificial diapause (AD), ovaries are surgically removed during the pre-implantation period to avoid E2 surge, followed by P4 injections to keep the uterus in the pre-receptive state and to allow embryo survival. It is postulated that the lack of ovaries, which is the source of nidatory E2, prevents embryo attachment (McLaren, 1971; Yoshinaga and Adams, 1966). The embryos survive in the uterus for several days and potentially weeks as long as P4 is present, and implantation can be induced by a single dose of E2 (Cha and Dey, 2014). Although E2 can induce attachment, there is additional evidence that if P4 is injected at the same time as removal of the ovary in the AD model, implantation can occur in the absence of E2 (McLaren, 1971; Yoshinaga and Adams, 1966).

Although the AD model is presumed to match the ND model, the two diapause models have not been compared for serum levels of E2 and P4 or for embryo location. We recently demonstrated that in a mouse natural virgin pregnancy (NP), embryos display three phases of movement: embryo entry, unidirectional clustered movement, and bidirectional scattering and spacing movement (Flores et al., 2020). Using the AD model it has been suggested that during diapause, embryos space out evenly but attachment does not occur (Nilsson, 1974). However, embryo location has not been assessed in lactating ND mice. In this study, we use these three models of pregnancy (**Fig. 1**), NP, ND, and AD, to assess serum levels of E2 and P4 and their impact on embryo location. We find that serum P4 levels in each model are vastly different, whereas serum E2 levels are relatively similar. In addition, we discovered that the ND and AD models of diapause differ in embryo location patterns. Further, unlike the NP model, the ND model displays embryo entry followed by a single unidirectional scattering and spacing movement. Finally, with the ND and AD models of pregnancy and exogenous hormone treatment of NP (**Fig. 1**), we show that both P4 and E2 can modulate different phases of embryo movement.

**Figure 1:**
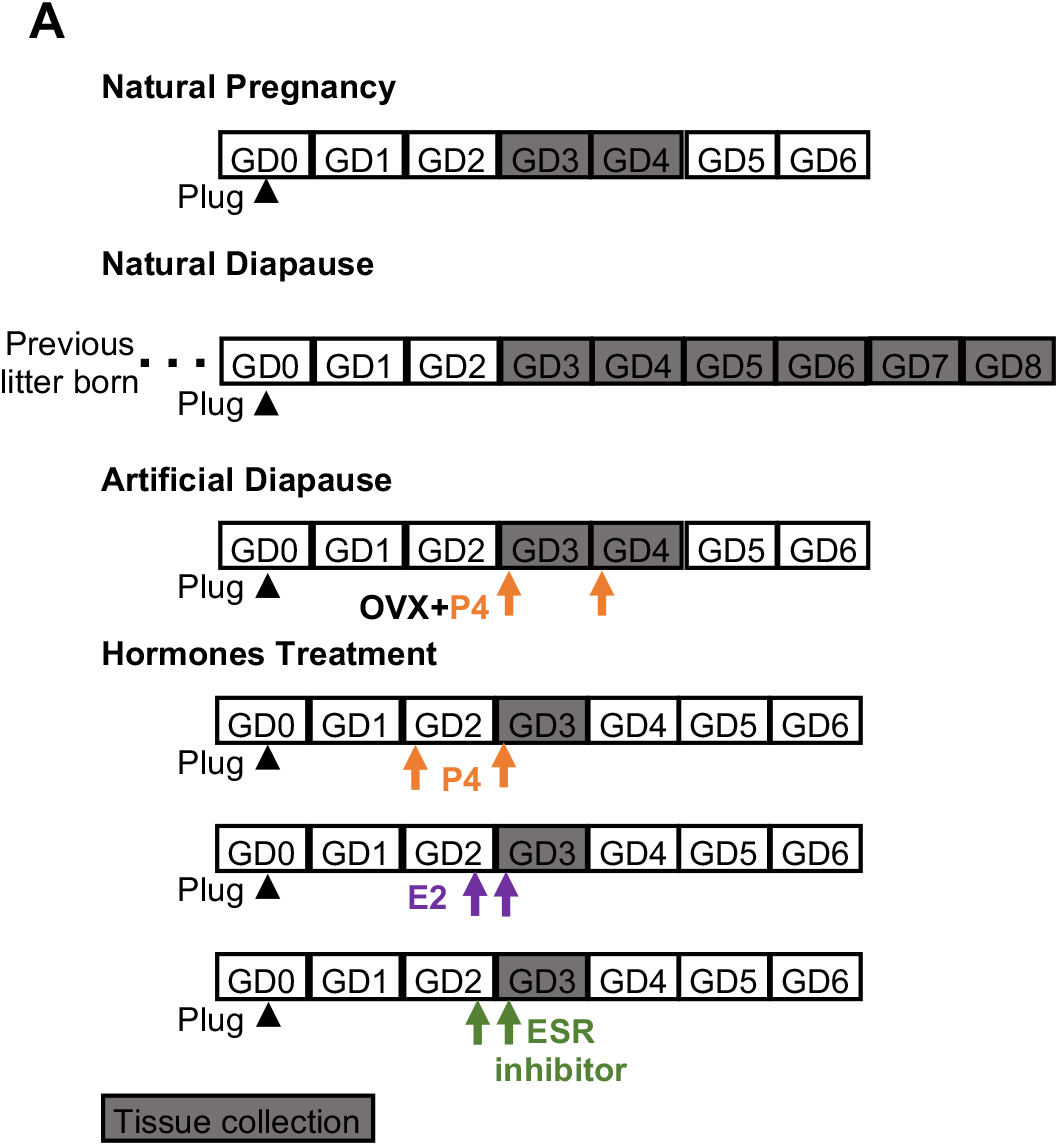
Schematic of models used to study the effects of ovarian hormone modulation on embryo location. Natural Pregnancy (NP) analysis is performed in the first pregnancy of a virgin post-pubertal mouse. For Natural lactational Diapause (ND), the female mouse gives birth to its first litter and mates with a male within 48 hours due to post-partum estrous, and the resulting pregnancy is analyzed. Artificial Diapause (AD) is a delay in implantation induced under laboratory conditions by removing the ovaries during the first pregnancy of a virgin mouse, and only P4 is injected to keep the pregnancy active. To corroborate results obtained from these models, we also use exogenous hormone (E2 or P4) administration or estrogen receptor (ESR) inhibitor on gestational day (GD) 2 or/and GD 3 to understand their effect on embryo movement. Arrowhead is G0.5 day of plug. Arrow indicates the injection time. OVX: ovariectomy.

## MATERIALS AND METHODS

### Animals

All animal research was carried out under the Michigan State University Institutional Animal Care and Use Committee guidelines. CD1 (ICR) mice aged 6 to 12 weeks were maintained on a 12 hours light/dark cycle. For Natural Pregnancy (NP) adult females were mated with fertile wild-type males to induce pregnancy. For Natural lactational diapause (ND) the female would carry pups to term and deliver. After delivery, female and male mice were in the same cages as the suckling pups. The appearance of a vaginal plug was identified as the gestational day (GD) 0.5 in all pregnancy models. Uterine dissections were performed at various time points from GD3 to GD8. For detecting implantation sites in ND on GD5 to GD8, 200 μl of 0.5% Evans blue dye (MP Biomedicals, ICN15110805) in phosphate-buffered saline (PBS) was injected into the lateral tail vein of the pregnant mouse 15 minutes prior to sacrificing the mouse. Uteri were then photographed in white light to observe implantation sites (Psychoyos, 1965).

### Artificial diapause procedure

For inducing artificial diapause, there are different methods proposed in the literature that differ in the timing of P4 treatment once ovaries are removed (McLaren, 1971; Yoshinaga and Adams, 1966) (Cha and Dey, 2014; McLaren, 1971; Paria et al., 1993). We chose the method that allows implantation to occur despite the removal of ovaries to minimize the variables in our study (McLaren, 1971; Yoshinaga and Adams, 1966). Thus, artificial diapause was induced by removing both ovaries on the morning of GD3 (0600h), leaving the oviduct intact. A subcutaneous injection of 2mg of P4 in 100μL of sesame oil was given at the time of surgery. Two groups of mice were dissected on GD3 at 1200h or 2200h. The third group of mice was injected with a second dose of 2mg P4 the following morning on GD4, and was dissected in the afternoon on GD4 (1500h - 1700h).

### Hormone and inhibitor treatments

17β-Estradiol (E8875, Sigma-Aldrich) and Progesterone (P0130, Sigma-Aldrich) were dissolved in sesame oil (Acros Organics, AC241002500) and injected subcutaneously at 25ng (Ma et al., 2003) and 4mg (Liang et al., 2018) respectively. Mice in the vehicle groups received injections of sesame oil alone. E2 was administered either on GD2 2200h or on GD3 0800h, while P4 was on GD2 at 1000h and GD3 at 1000h. E2 inhibitor ICI 182,780 (104710, Tocris Bioscience) was dissolved in DMSO, and the stock solution was diluted in sesame oil, and 50μg was injected subcutaneously (Cha et al., 2013). Mice in the vehicle group received subcutaneous injections of 0.1mL sesame oil. E2 inhibitor or vehicle was administered on GD2 2200h, and mice were dissected at GD3 1500h. An additional group of mice was injected at GD2 2200h and a second injection at GD3 1000h and dissected at GD3 2200h.

### Blood serum collection

At the time of dissection, mouse blood was collected by cardiac puncture. Serum isolation protocol was modified from the University of Virginia Ligand Assay and Analysis Core. Briefly, blood was allowed to clot for ~30 minutes. Then a plastic pipette tip was used to disrupt the clot before centrifuging at 2000Xg for 15 minutes. Serum was separated from the rest of the blood cells and stored in an Eppendorf tube at −20°C. E2 and P4 levels were assayed by the University of Virginia Center for Research in Reproduction Ligand Core.

### Whole-mount immunofluorescence

Whole-mount immunofluorescence staining was performed as described previously (Arora et al., 2016). Uteri were fixed in DMSO:Methanol (1:4) after dissection. To stain the uteri, they were rehydrated for 15 minutes in 1:1, Methanol: PBST (PBS, 1% Triton X-100) solution, followed by a 15 minutes wash in 100% PBST solution before incubation. Samples were incubated with Hoechst (Sigma Aldrich B2261) diluted in PBST (1:500) for two nights at 4°C. The uteri were then washed once for 15 minutes and three times for 45 minutes each, using PBST. Next, the uteri were stretched in 100% methanol, followed by 30 minutes dehydration in 100% methanol, an overnight incubation in 3% H_2_O_2_ solution diluted in methanol, and a final dehydration step for 60 minutes in 100% methanol. Finally, samples were cleared using a 1:2 mixture of Benzyl Alcohol: Benzyl Benzoate (Sigma 108006, B6630).

### Confocal microscopy

Confocal imagining procedures were done as previously described in Flores *et* al. Stained uteri were imaged using a Leica TCS SP8 X Confocal Laser Scanning Microscope System with white-light laser, using a 10x air objective. For each uterine horn, z-stacks were generated with a 7.0 μm increment, and tiled scans were set up to image the entire length and depth of the uterine horn (Arora et al., 2016). Images were merged using Leica software LASX version 3.5.5.

### Image analysis for embryo location

Image analysis was done using commercial software Imaris v9.2.1 (Bitplane). Embryo location was assessed as described previously(Flores et al., 2020). Briefly, confocal LIF files were imported into the Surpass mode of Imaris and using the Surface module 3D renderings were used to create structures for the oviductal-uterine junctions, embryos, and horns. The three-dimensional Cartesian coordinates of each surface’s center were identified and stored using the measurement module. The distance between the oviductal-uterine junction and an embryo (OE), the distance between adjacent embryos (EE), and the horn length was calculated using the orthogonal projection onto the XY plane. All distances were normalized to the length of their respectively uterine horn Horns with less than three embryos were excluded from the analysis. These distances were used to map the location of the embryos relative to the length of the uterine horn. The uterine horn was divided into three equally spaced segments –closest to the oviduct, middle, and closest to the cervix. These segments were quantified for the percentage of embryos present in each section. Embryos in the oviductal region close to the oviductal-uterine junction were accounted for in the first oviductal segment.

### Statistical Analysis

For serum hormone levels, the unpaired two-tailed t-test was performed with Welch’s correction, while for ND and AD models, ANOVA was used to analyze OE and EE distances amongst uterine horns and different time points as described previously (Flores et al., 2020). Statistical analyses were performed with Graph Pad Prism. To compare the vehicle and the hormone or inhibitor treatments, linear mixed-effects models (Laird and Ware, 1982) were conducted to study the effect of treatment, time, and their interaction on OE and EE distances, while controlling for repeated measures within a horn using a random intercept term. Multiple comparisons of means were conducted using Tukey contrasts with the Holm method adjustment for p-values. Analysis was conducted using Graph Pad Prism, with advanced statistical analysis conducted using R Statistical Software (Team, 2014) and the nlme package (Pinehiro et al., 2021).

## RESULTS

### Ovarian hormones in a natural pregnancy

We first measured serum levels of ovarian hormones P4 and E2 in the NP model. We found that P4 levels are low on gestational day (GD) 0 and GD1, begin to rise on GD2 and remain high until implantation (GD4 0000h) (**Fig. 2**). On the other hand, serum E2 levels stayed constant throughout the pre-implantation phase of pregnancy, and we did not observe a nidatory rise in the serum levels of E2 (**Fig. 2**).

**Figure 2:**
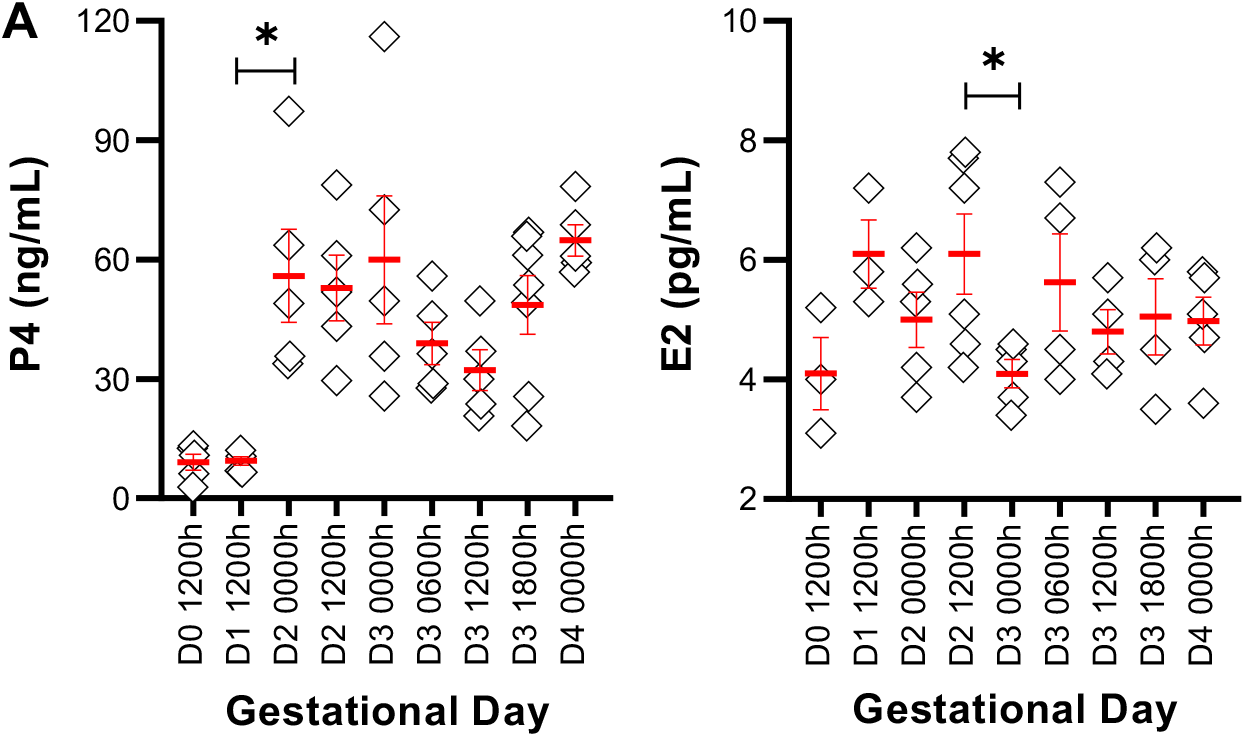
Ovarian hormones in a natural pregnancy. P4 and E2 were measured in mouse serum from GD0 to GD4. P4 levels begin to rise on GD2 (0000h), whereas serum E2 levels remain similar throughout early pregnancy. Each diamond represents data from one mouse. Mean ± SEM displayed in red. * indicates p<0.05.

### ND model displays two pauses separated by unidirectional embryo movement

In the NP model, on GD3 between 0600h and 1200h, 100% of the embryos remain clustered in the oviduct (**Fig. 3A**). At GD3 2100h ~88% of the embryos are still near the oviductal-uterine junction, with only 12% of the embryos displaying a displacement away from the oviduct. Thus, for most of GD3 embryo location is closer to the oviduct in the ND model. We compared the distribution of the embryos in the three segments of the uterus (near the oviduct - oviductal, middle, and near the cervix – cervical) and observed that on GD4, embryos appear to scatter and space unidirectionally throughout the length of the horn. At GD4 0900h, 60% of embryos are present in the oviductal segment of the horn, 28% in the middle, and 2% in the cervical segment, while by the evening of GD4 (1800h), 39% of embryos are in the oviductal segment, 35% in the middle, and 26% in the cervical segment (**Fig. 3A**). The percentage of embryos steadily increases in the middle and cervical segments until an equal distribution is achieved on GD6 (**Fig. 3A**). These data suggest that embryo clusters in the ND model are paused near the oviduct-uterine junction on GD3, followed by robust embryo movement on GD4, fine embryo spacing by GD6 followed by a second pause until the embryos attach (**Fig. 3A**).

**Figure 3:**
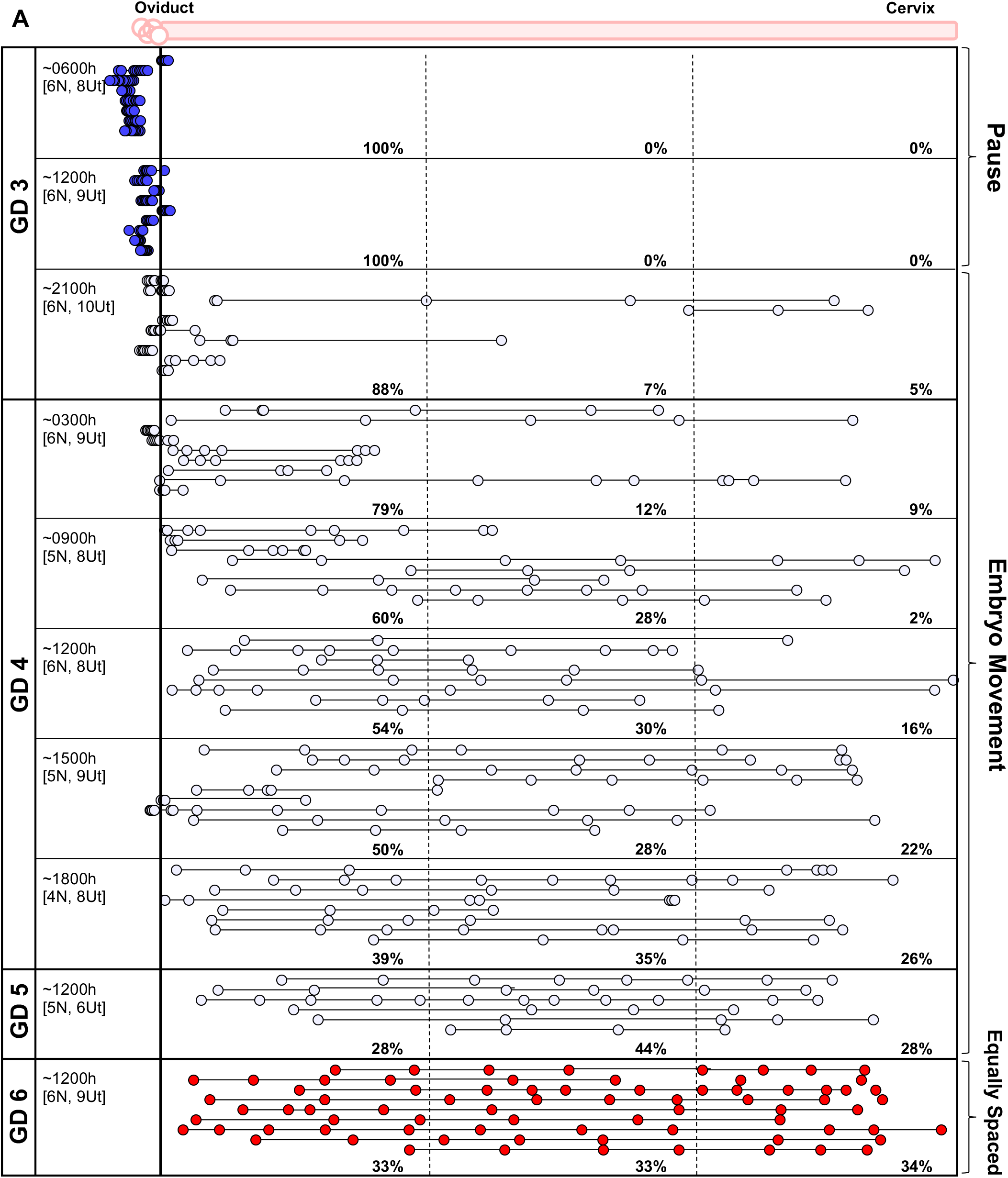
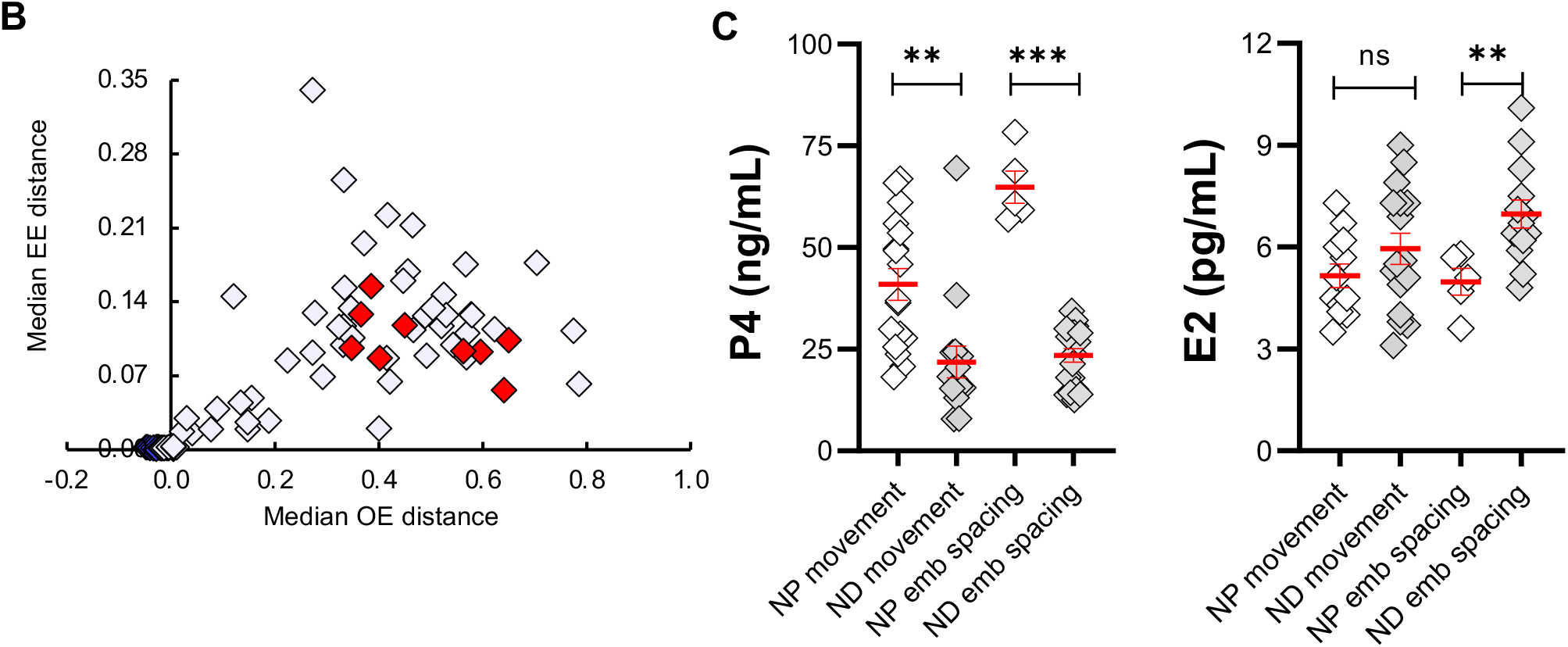
Embryo movement analysis in a lactational natural diapause pregnancy. **(A)** Location of embryos in uterine horns at fixed time intervals on GD3 – GD6 in ND. Each circle represents an embryo, and circles connected with a line are embryos from the same uterine horn. Blue-white-red colors signify time scale where blue represents time points on GD3 when embryos are at the oviductal-uterine junction, white represents embryo movement on GD3, GD4, and GD5, and red represents timing when embryos are equally spaced out and attachment is first observed in some ND pregnancies. The left-hand column indicates the time of dissection in hours (h). ‘N’ represents the number of mice, and ‘Ut’ represents the number of uterine horns analyzed for each time point. Dotted lines divide the uterine horns into three equal segments, and percentages for each time point signify the percentage of embryos in each segment. **(B)** OE vs. EE analysis of embryo location data from (A). Each diamond represents the median value from individual uterine horn from (A) and is color-matched for GD. **(C)** Comparison of serum P4 and E2 in NP and ND. Time points are combined, and hormone levels are compared during the movement phase or after embryo spacing. P4 levels are lower both during embryo movement and attachment. E2 levels in ND pregnancy are slightly higher than NP. Mean ± SEM displayed in red. ns: not significant, ** indicates p<0.01, *** indicates p<0.001.

We examined the median oviduct-embryo (OE) distance and median embryo-embryo (EE) distance per horn (Flores et al., 2020) throughout embryo movement in ND. In the initial stages, the embryos display smaller OE and EE distances (**Fig. 3B**), supporting the presence of embryo clusters near the oviduct. As the embryos move, the OE and the EE distances increase simultaneously until equal spacing is achieved (**Fig. 3B**). These data further support a unidirectional scattering and spacing embryo movement pattern towards the cervix (Boving, 1956) in ND.

### Implantation reaction in ND model

Control mice display a blue dye reaction indicating vascular permeability on the evening of GD4, confirming implantation (Restall and Bindon, 1971). However, when the blue dye was administered to ND mice, as expected, we did not see a blue dye reaction on GD4 (**data not shown**) and GD5 (**Fig. S1**), suggesting that implantation initiates after these time points. The first signs of blue dye reaction were observed on GD6, where 40% of the mice displayed a positive blue dye reaction (**Fig. S1**). This suggests that there is at minimum a two-day delay in attachment initiation, which is in line with the minimum observed delay in delivery (Mantalenakis and Ketchel, 1966; Norris and Adams, 1981). At GD7, 43%, and on GD8, 89% of the ND mice displayed a positive blue dye reaction (**Fig. S1**).

### Serum P4 levels but not E2 levels are different between NP and ND

To compare ovarian hormone levels across NP and ND models, we evaluated serum P4 and E2 levels during embryo movement and after embryos achieved equidistant spacing. For NP, values for time points for embryo movement ranging from GD3 0600h – GD3 1800h (Flores et al., 2020) were pooled, and for post embryo spacing, GD4 0000h was used. For ND, values for time points for embryo movement ranging from GD3 2200h – GD5 1200h were pooled, and for post embryo spacing time points from GD6 1200h – GD8 1200h were pooled. Surprisingly, we found that the levels of serum P4 in ND were lower than NP both during embryo movement (mean P4 in NP 40.95 ng/ml, in ND 21.85 ng/ml, p<0.01), and after embryo spacing (mean P4 in NP 64.82 ng/ml, in ND 25.05 ng/ml, p<0.0001) (**Fig. 3C, Fig. S2**). In addition, although E2 levels were also different between NP and ND post embryo spacing, the trend was towards higher E2 levels in ND (mean: 6.97 pg/ml) compared to NP (mean: 4.98 pg/ml, p<0.01) (**Fig. 3C, Fig. S2**).

### Exogenous E2 affects both unidirectional and bidirectional embryo movement

In addition to differences in ovarian hormones, the uterine milieu in the ND model is recovering from the prior pregnancy, and is likely affected by wound repair factors and the presence of immune cells. To distinguish between the effects of ovarian hormones from other changes in the uterine milieu, we tested the specific effects of E2 and P4 on embryo location by exogenous hormone treatment in NP. E2 and P4 exogenous treatments were performed at times when endogenous hormones are normally present. This was done to ensure that hormone receptor expression in appropriate compartments would be present and exogenous treatment would merely amplify the hormone receptor signaling.

We treated NP mice with 25ng E2 subcutaneously. One set of mice was treated with E2 on GD2 (2200h) prior to embryo entry to assess the effect on unidirectional clustered movement at GD3 1200h. At GD3 1200h, in both vehicle and E2 treatment, embryos were observed in clusters. However, embryos accumulated primarily in the middle segment upon E2 treatment compared to the first segment for vehicle treatment (**Fig. 4A**). Both the mean OE and EE distance for E2 treatment was larger than vehicle treatment (p<0.005). This suggests that E2 may increase the velocity of embryo movement in the unidirectional phase. Another set of mice was treated with E2 on the morning of GD3 (0800h) to assess effects on bidirectional movement (GD3 1500h and 2200h). At GD3 1500h, the majority of the embryos were present as clusters in the middle segment of the uterine horn in vehicle and treatment (51% and 42% respectively, **Fig. 4B**). On the other hand, when evaluated later in the day (GD3 2200h), more embryos were present in the oviductal segment (52%) and fewer embryos were found in the cervical segment (12%) as compared to a nearly equal distribution of embryos in all three segments in the vehicle-treated animals (**Fig. 4C**). Further while the mean EE distance was similar in vehicle and E2 treatment, the mean OE distance was significantly different (p<0.05) supporting altered embryo distribution along the uterine horn.

**Figure 4:**
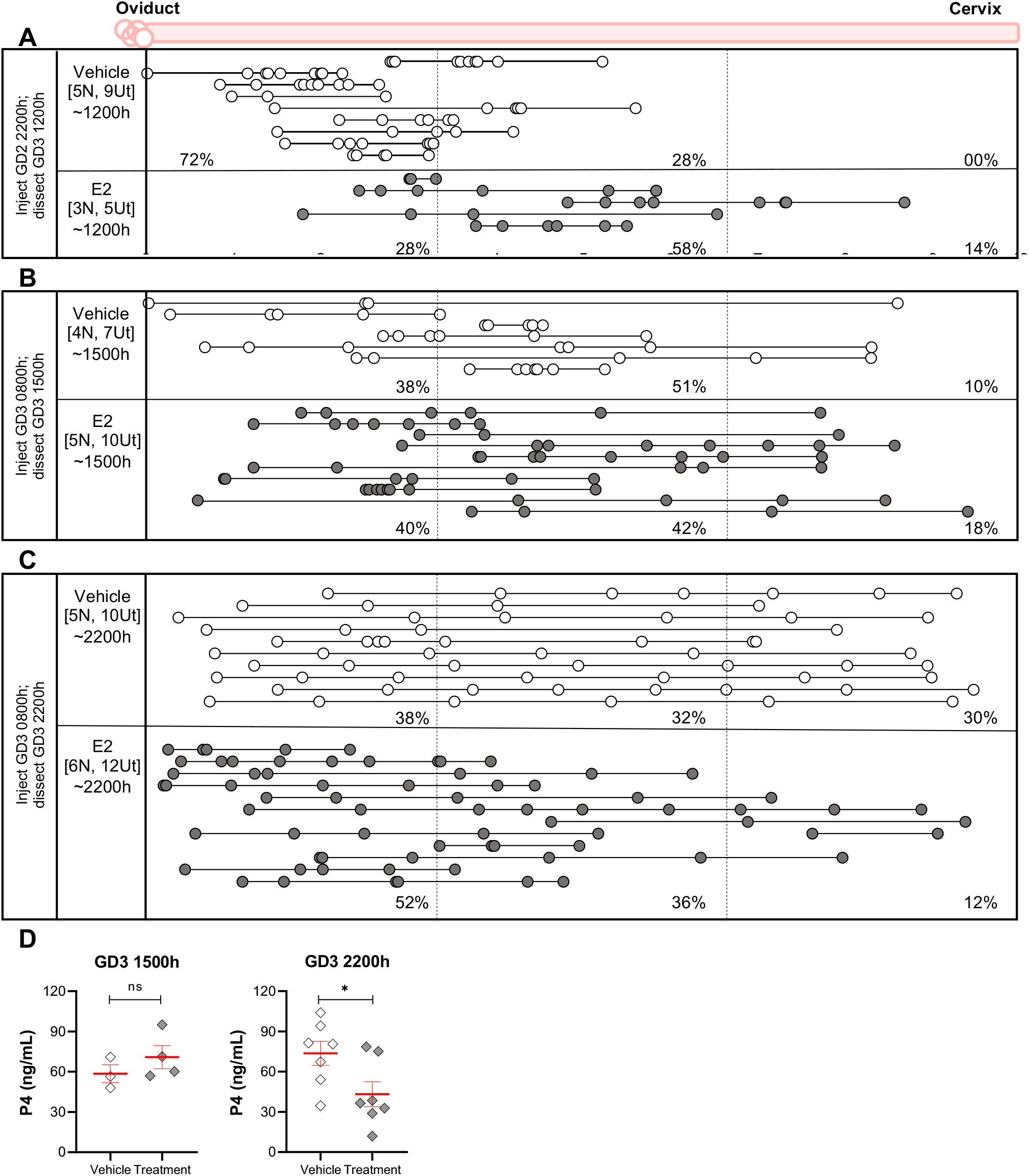
E2 treatment displays an immediate effect on unidirectional and delayed effect on bidirectional embryo location. Treating the NP with E2 prior to embryo entry alters the embryo movement pattern in the unidirectional phase (A). Treatment with E2 after embryo entry does not alter embryo location at the beginning of the bidirectional phase (B), but there is a delayed impact on embryo location in the latter half of the bidirectional phase (C). (D) P4 levels were measured after E2 treatment and were similar in the first half of bidirectional movement (GD3 1500h) but were reduced at 2200h. Mean ± SEM displayed in red. ns: not significant, * indicates p<0.05.

To address the altered embryo location with E2 treatment, we evaluated P4 levels in vehicle and E2 treated mice. While P4 levels were comparable between vehicle and treatment at GD3 1500h, mean values 58.52 and 70.86 ng/ml respectively, P4 levels at GD3 2200h were higher in vehicle (mean 73.7 ng/ml) and reduced after E2 treatment (mean 43.19 ng/ml, p<0.05) (**Fig. 4D**). These data suggest a delayed impact of E2 treatment on the bidirectional phase of embryo movement, possibly through a reduction in P4 signaling.

### Inhibiting E2 activity does not affect unidirectional or bidirectional embryo movement

We then tested the effect of inhibiting E2 activity on embryo location patterns. We treated NP mice with a commonly used estrogen receptor (ESR) inhibitor ICI 182,780on GD2 2200h and GD3 0800h and evaluated embryo location at GD3 (1500h) and GD3 (2200h). Inhibiting E2 activity by blocking ESR did not impact embryo location in the NP model of pregnancy (Fig. S3).

### P4 treatment affects the unidirectional phase of embryo movement

To assess the effects of P4, we treated females mated for the NP with 4mg P4 subcutaneously (**Fig. 5**). First, to address the effect of P4 treatment on embryo entry, we treated mice with P4 on GD2 1000h and assessed embryo location on GD3 0300h. Embryo location was equivalent in both vehicle and P4 treatment, suggesting an identical pattern for embryo entry (**Fig. 5A**). Next, we treated mice with P4 on GD2 1000h and GD3 1000h and evaluated them at GD3 1500h or 2200h. On GD3 1500h, in the P4 treatment group, most of the embryos (72%) were present near the oviductal-uterine junction and in the oviductal segment of the uterus, compared to vehicle-treated controls. Both mean OE and EE distances were significantly different between vehicle and P4 treated mice (p<0.0001 and p<0.005 respectively). This embryo location was reminiscent of a unidirectional scattering and spacing embryo movement (**Fig. 5B**). When uteri were evaluated later in the day at GD3 2200h, the embryos appeared to space out evenly, similar to the vehicle-treated controls (**Fig. 5C**), suggesting that P4 affects the earlier unidirectional clustered movement, but the embryos space out eventually. We postulated that the loss of effect on embryo movement might be due to a drop in P4 levels later in the day. Although there was a reduction in serum P4 levels at GD3 2200h (mean value 414.8 ng/ml), compared to GD3 1500h (values >800 ng/ml), these levels were much higher compared to vehicle-treated mice (mean value 48.02 ng/ml at GD3 1500h and mean value 75.83 ng/ml at GD3 2200h) (**Fig. 5D**). E2 levels remained similar in treatment vs. vehicle at both time points (Fig. 5E).

**Figure 5:**
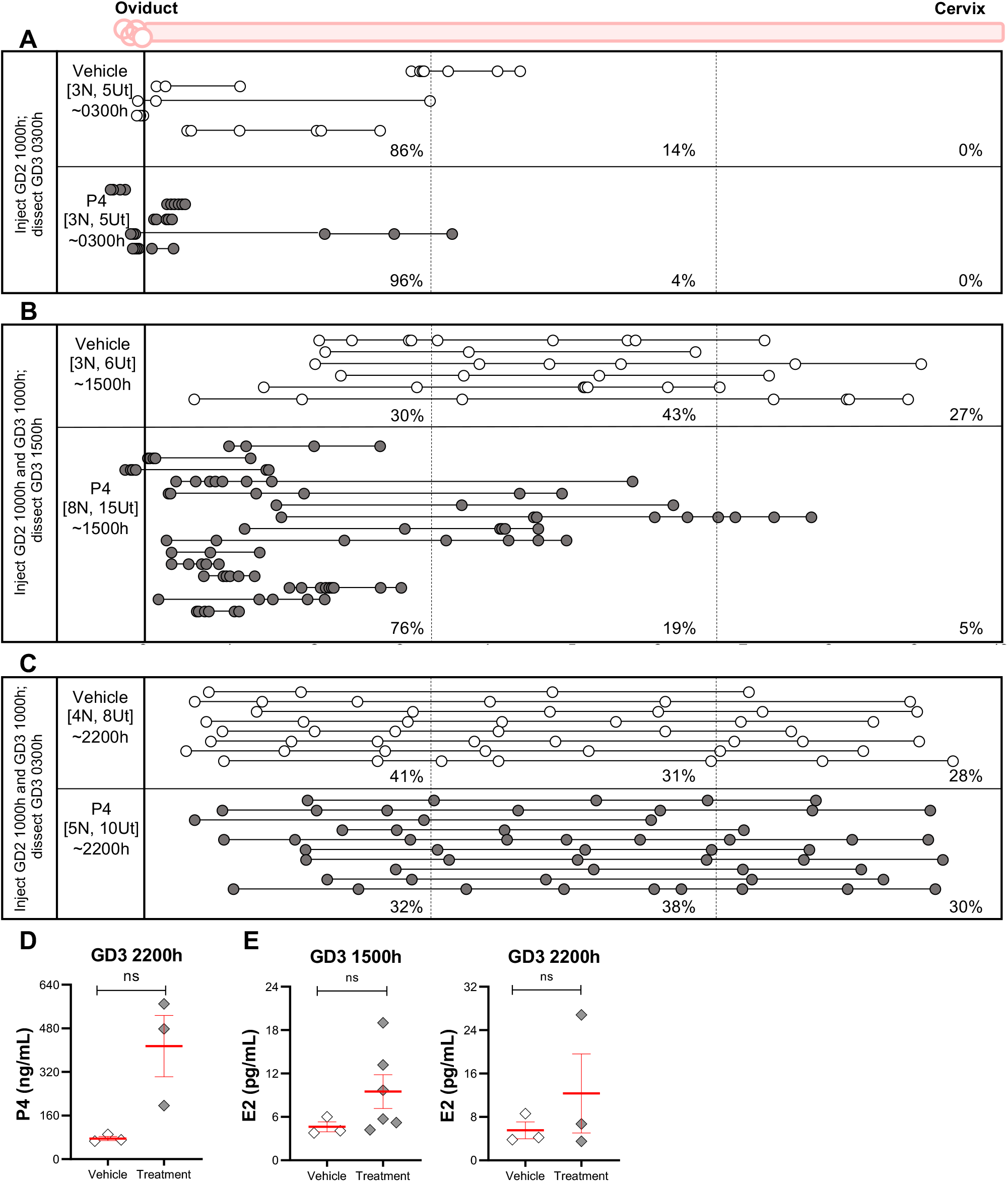
P4 treatment displays an immediate effect on embryo location that resolves over time. Treating the NP with P4 does not affect embryo entry (A). P4 treatment for NP induces embryo location concordant with a slower unidirectional embryo movement pattern on GD3 1500h (B). (C) Although the embryos initially display an altered movement pattern, they eventually space out at the end of bidirectional movement on GD3 2200h. (D) P4 levels and (E) E2 levels upon P4 treatment. Mean ± SEM displayed in red. ns: not significant.

### Artificial diapause mimics Natural Pregnancy in embryo location patterns

We evaluated embryo location in the AD model of pregnancy that represents a laboratory-induced delay in embryo attachment. AD was induced on the morning of GD3, and embryo location was evaluated in these mice at different time points. On GD3 1200h, we observed that embryos are primarily found in the middle segment of the uterine horn, with 26% of embryos in the oviductal segment, 53% in the middle segment, and 21% in the cervical segment. Further, when evaluated at GD3 2200h, the embryos largely remained in the middle segment (**Fig. 6A**). The embryos had achieved even spacing by GD4 mid-day (1500h – 1700h) (**Fig. 6B**), suggesting that embryo spacing occurs normally in the AD model.

**Figure 6:**
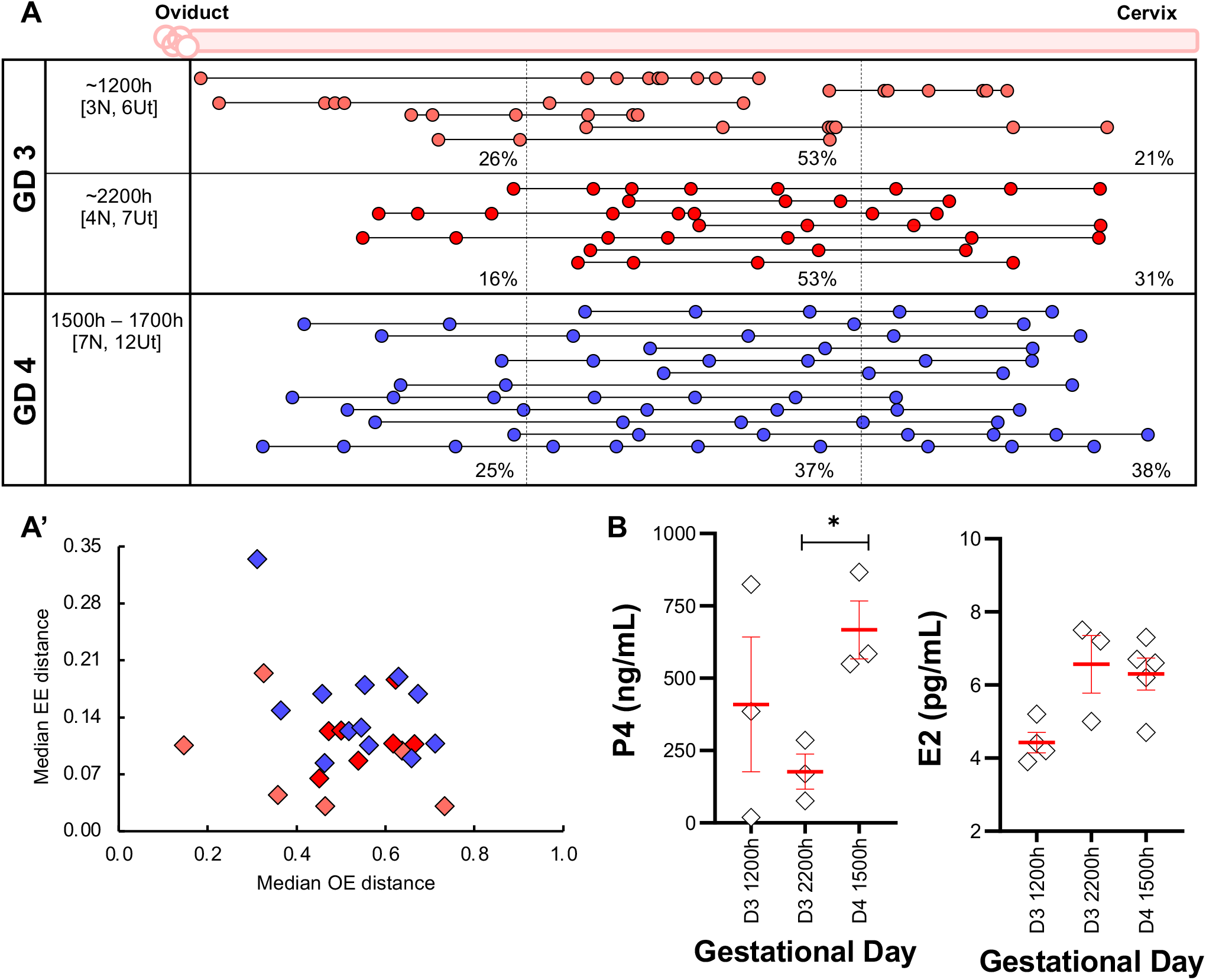
Embryo movement analysis in artificially induced diapause pregnancy. **(A)** Ovariectomy and treatment with P4 on the morning of GD3 displays embryo location consistent with unidirectional clustered movement (A). On GD4, embryos eventually space out despite higher levels of P4. (**A’**) OE and EE distribution of uterine horns from (A). **(B)** Serum P4 levels after 2mg P4 doze are much higher than NP, and serum E2 levels remain constant despite ovariectomy suggesting an alternate source of E2. Mean ± SEM displayed in red. * indicates p<0.05.

### Serum P4 levels but not E2 levels are different between NP and AD

We further evaluated the serum P4 and E2 levels in the AD model during embryo movement and compared them with NP and ND models of pregnancy. We found that a 2mg dose of P4 in the AD model induced ~10-fold high P4 concentration (~385.40 ng/ml) (**Fig. 6C**) compared to the NP model during embryo movement (mean 40.95 ng/ml) and about 20-fold those found of the P4 levels found in the ND model during embryo movement (mean 21.85 ng/ml). In contrast, serum E2 levels appeared to be similar across NP, ND, and AD in the embryo movement period, showing mean E2 values of 5.16 pg/ml, 5.95 pg/ml, and 5.34 pg/ml, respectively. Thus, E2 levels in the AD model point to an alternate source of E2 in the ovariectomized mice, and contrary to the prevalent notion, P4 and not E2 is different between all three models of pregnancy.

## DISCUSSION

Pausing pregnancy due to unfavorable birthing conditions is an evolutionary mechanism to ensure the survival of the young. Hormonal regulation of the uterine milieu plays a key role in ensuring survival and pregnancy success when the conditions do become favorable. Reproducing such paused state in the laboratory conditions allows reproductive and developmental biologists to ask questions pertaining to hormonally altered mechanisms that support pregnancy survival. We utilized different mouse models of pregnancy to understand variation in ovarian hormone levels and their possible effects on embryo movement patterns during pre-implantation stages of the pregnancy.

### Differences between ND and NP model and two pauses in ND

We observed that embryo location and movement patterns differed in the first virgin pregnancy (NP) and the lactational diapause pregnancy (ND). Embryos in NP are found near the oviductal uterine junction on the morning of GD3 and undergo clustered, scattering, and spacing movements on day 3 of pregnancy before attaching at GD4 (Flores et al., 2020). On the other hand, although similar to NP, embryos in the ND model of pregnancy arrive at the oviductal-uterine junction on GD3, unlike NP, they stay near the junction for an entire day. This suggests that in the ND model there is a 24-hour (first) pause around the time of embryo entry into the uterus but prior to embryo movement. This first pause in the ND mouse model resembles the diapausing mink, where blastocysts are clustered near the anterior portion of the uterus (Fenelon et al., 2014) and the diapausing armadillo where blastocysts are in the portion of the uterus closer to the oviductal opening (Enders and Buchanan, 1959). In the mink, embryo spacing occurs once the pregnancy is reactivated. However, we noted that in the ND mouse model, the embryos enter the uterus and begin to scatter after the first 24-hour pause. Unlike the unidirectional clustered movement followed by the bidirectional scattering embryo movement in the NP model (Flores et al., 2020), embryo movement in the ND model is consistent with a unidirectional scattering and spacing pattern. Embryo scattering begins on GD4, but even spacing is slowly acquired by GD6, followed by a second pause until embryo attachment occurs. This pattern of movement and spacing is reminiscent of embryo movement in the rabbit’s natural pregnancy (Boving, 1956).

While the earliest signs of attachment in the NP model are observed at GD3 2100h (Restall and Bindon, 1971), the earliest attachment in ND was observed at GD6, indicating a minimum 2-day delay. This is consistent with the 24-hour pause followed by slow embryo movement in ND. These data are consistent with mouse breeding observation where first litters are observed at ~19-20 days, whereas second litters are observed ~21-22 days after the first litter of pups is born (Mantalenakis and Ketchel, 1966; Norris and Adams, 1981). The number of suckling pups affects the time delay in attachment in the ND model (McLaren, 1968; Norris and Adams, 1981). We did not normalize the number of pups for our ND pregnancies, thus a variable number of pups could explain differing timing of implantation in our ND model (Mantalenakis and Ketchel, 1966).

### Similarities between AD and NP model and one pause in AD

The AD model is derived from the NP model, and although it is assumed to mimic the ND model, we found that the AD model displays embryo location parameters similar to the NP model. In both the NP and AD models, embryos move as clusters to accumulate in the center of the horn before bidirectionally scattering and spacing out (Flores et al., 2020). AD mice are generated by surgically removing the ovaries on the morning of GD3 when embryos have already entered the uterine horn as clusters (Flores et al., 2020). The removal of the ovaries and exogenous supplementation of P4 in AD permits embryo spacing consistent with previous data (Nilsson, 1974). Thus, the AD pregnancy displays only a single pause after embryo spacing but prior to embryo attachment.

### Serum P4 levels differ across NP, ND, and AD

Implantation in NP is thought to result from the nidatory peak of E2 (Ma et al., 2003). Our analysis of the different models of pregnancy suggests that serum E2 levels tend to stay basal across all models during the embryo entry, embryo movement, and paused states before embryo attachment. The lack of variation in observed E2 levels in our study could be due to the sensitivity of the assay to detect differences in the serum E2 levels. Alternatively, E2 levels in the serum may stay basal, and there may be a direct route of delivery of E2 through the ovarian circulation into the uterus (Tourgeman et al., 2001). However, to-date there is no such report in mice.

In contrast to E2, P4 levels in ND are roughly half that of NP, and in AD are ~10 times that of NP. Thus, in the ND model, lower P4 and not E2 may be the reason for altered unidirectional movement of embryos and also for the delay in implantation. The latter observation is also supported by studies where similar to exogenous E2, exogenous P4 injections can also support implantation both in the ND rat model (Yoshinaga, 1961) and in the ND and AD mouse model (McLaren, 1971). As part of post-partum endometrial regeneration (Yoshii et al., 2014), ND model must be characterized by wound healing responses, including secretion of prostaglandins and cytokines in the uterus to repair placental scars (Mackler et al., 2000). These factors and the scars may impact embryo movement in the ND model and are not accounted for in our study. Nevertheless, similar levels of serum E2 across the different pregnancy models with varying embryo movement patterns suggest that P4 may primarily regulate embryo movement in the uterus.

### AD and ND models: using caution while comparing results

AD has been used as a model to study delayed implantation for decades (Cha and Dey, 2014; McLaren, 1971; Paria et al., 1993). Due to the ease of generating AD animals and cost-effectiveness, it has been used as a proxy to understand the transcriptional, proteomic, and structural changes in the embryo or the uterus that support implantation (Fu et al., 2014; Hamatani et al., 2004; He et al., 2019; McLaren, 1971; Nilsson, 1974). However, when comparing embryo location and hormone levels between ND and AD, we noted stark differences. The ND model displays two pauses, whereas the AD model only represents a single pause. Thus, the AD model cannot be used to study the state of the embryo or the uterus in the first pause in the ND model. Even after embryos are spaced out, serum P4 levels in the ND model are much lower than aberrantly high P4 levels in the AD model. Thus, the AD model can still be used to understand aspects of delayed implantation (ND), however, it is important to recognize the differences between the two models and exercise caution when interpreting the results.

### E2 and P4 influence embryo movement patterns

Ovarian hormone levels are crucial for a receptive uterus and for uterine muscle (myometrial) contractions. In women undergoing in vitro fertilization (IVF), uterine contraction frequency is reduced when higher P4 levels are present on the day of embryo transfer (ET). Further, ovulatory E2 promotes uterine contractions, but E2 treatment in the presence of P4 may not affect uterine contractility (Bulletti and de Ziegler, 2006; de Ziegler et al., 2001; Sebag-Peyrelevade and Fanchin, 2015). Hormonal regulation of uterine contractility could directly impact embryo movement. In our study, blocking E2 signaling by using an ESR inhibitor did not affect embryo movement in the unidirectional or bidirectional phase of the NP model. For exogenous hormone treatment, E2 accelerated while P4 slowed the clustered embryo movement after entry into the uterus, and these effects could be attributed to ovarian hormone modulation of uterine contractility. Similar to contractions in women undergoing IVF and ET, higher P4 levels upon exogenous P4 treatment correlate with the initial slow embryo movement, although embryos do space out evenly eventually. This data is reminiscent of our recently published work where modulating muscle contractions in the unidirectional clustered phase prevents embryo movement. On the other hand, modulating P4 (this study) and uterine contractions (Flores et al., 2020) in the bidirectional phase does not affect embryo movement. These observations lead us to hypothesize that P4 and E2 may regulate embryo movement by modulating uterine contractions during early pregnancy.

The bidirectional phase of movement appears to be dependent on embryo-uterine interactions (Flores et al., 2020). Our study showed a delayed effect of E2 treatment on the bidirectional phase of embryo movement. Since relaxing the muscle does not affect the bidirectional phase of embryo movement, the delayed effect of E2 could be due to embryo-uterine signaling triggered due to transcriptional activity of E2. Alternatively, P4 levels were reduced in E2 treated mice and could signal either a loss of pregnancy or mimic a state of diapause (McLaren, 1971) and account for altered embryo movement patterns. We were unable to directly address the impact of reduced P4 signaling using an inhibitor of progesterone receptor such as Mifepristone because such treatment affects embryo viability (Roblero et al., 1987). In order to further confirm whether E2 and P4 regulate embryo movement by modulating muscle contractions in the first phase of movement and alternate mechanisms in the second phase of movement, hormone receptors should be depleted in the uterine smooth muscle compartment using appropriate Cre driver lines (Ghosh et al., 2020). This will be the subject of future studies.

In conclusion, we show that ND, AD, and NP are distinct models of pregnancy with variable serum P4 but similar E2 levels. Embryo movement patterns are distinct between the first virgin pregnancy (NP) and lactational diapause (ND), and also when exogenous hormones are administered, implicating at least partial hormonal regulation of embryo movement in the uterus.

These results have clinical significance in ovarian hyper-stimulation and artificial reproductive technologies, where higher P4 levels during blastocyst transfer could impact embryo movement, affecting embryo location during implantation and eventually pregnancy outcomes.

## ACKNOWLEDGMENTS

We thank Asgerally Fazleabas and Gregory Burns for scientific discussions for the manuscript.

## COMPETING INTERESTS

The authors declare no competing interests.

## AUTHOR CONTRIBUTIONS

H.L, D.F. and R.A. designed the experiments; H.L, D.F., M.M., and Z.R. performed experiments; H.L, D.F., Z.R., A.C., M.D., and R.A. analyzed the data. H.L., D.F., and R.A. interpreted the results; H.L., D.F., and R.A. wrote the manuscript.

## FUNDING

We acknowledge support from March of Dimes grant #5-FY20-209

---

**Supplementary Figure 1:**
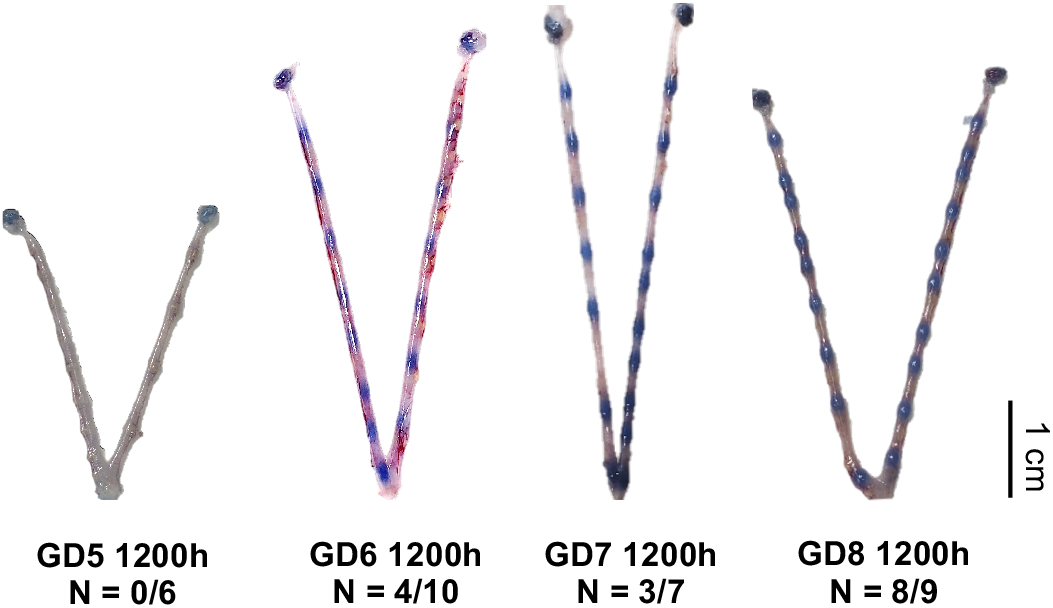
Implantation frequency in ND. Implantation begins on GD6 (4/10), and most ND pregnancies display implantation by GD8 (8/9) as seen by blue dye reaction. Bulges observed at GD5 are placental scars from prior pregnancy.

**Supplementary Figure 2:**
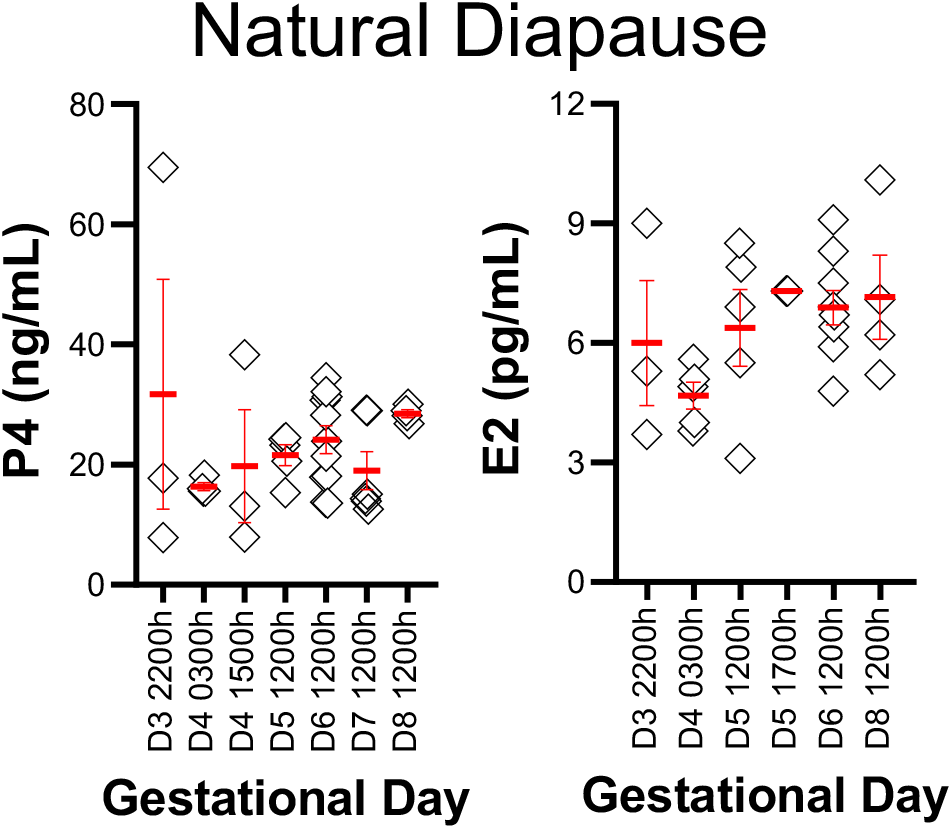
Serum ovarian hormones in ND pregnancy. Serum hormone levels during different time points of embryo entry (GD3), embryo movement (GD4-GD5), and post-embryo spacing (GD6) in the ND pregnancy. Mean ± SEM displayed in red.

**Supplementary Figure 3:**
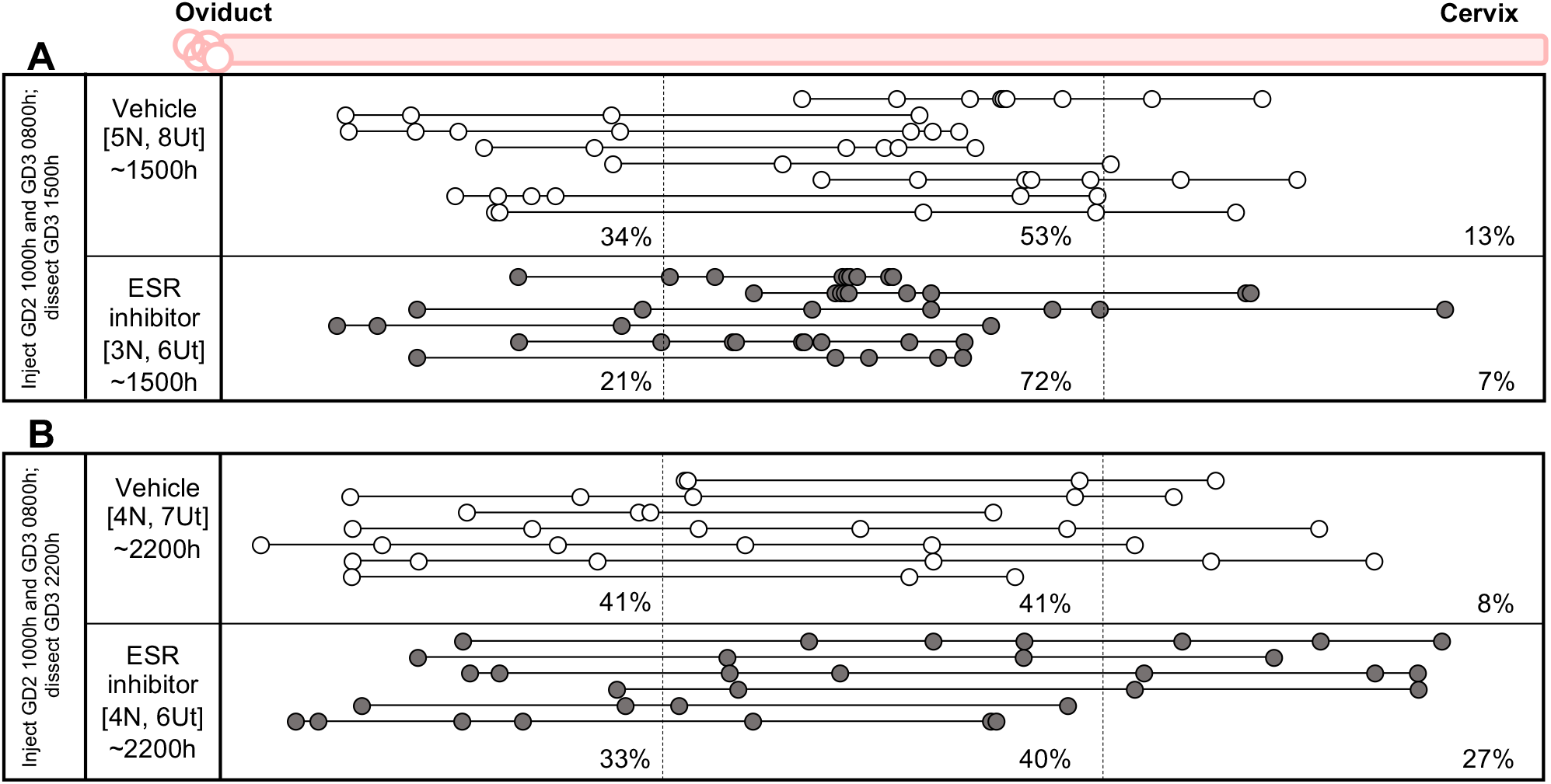
ESR inhibitor treatment does not affect embryo location. Treating the NP with ESR antagonist during unidirectional movement (A), or bidirectional movement (B), fails to alter the movement pattern of embryos, suggesting no effect of E2 levels on embryo movement.

## Notes

### Competing Interest Statement

The authors have declared no competing interest.

## REFERENCES

Arora, R., Fries, A., Oelerich, K., Marchuk, K., Sabeur, K., Giudice, L. C. and Laird, D. J. (2016). Insights from imaging the implanting embryo and the uterine environment in three dimensions. Development 143, 4749–4754.

Boving, B. G. (1956). Rabbit blastocyst distribution. The American journal of anatomy 98, 403–434.

Bulletti, C. and de Ziegler, D. (2006). Uterine contractility and embryo implantation. Current opinion in obstetrics & gynecology 18, 473–484.

Bylander, A., Gunnarsson, L., Shao, R., Billig, H. and Larsson, D. G. (2015). Progesterone-mediated effects on gene expression and oocyte-cumulus complex transport in the mouse fallopian tube. Reproductive biology and endocrinology : RB&E 13, 40.

Cha, J. and Dey, S. K. (2014). Cadence of procreation: orchestrating embryo-uterine interactions. Semin Cell Dev Biol 34, 56–64.

Cha, J., Sun, X., Bartos, A., Fenelon, J., Lefèvre, P., Daikoku, T., Shaw, G., Maxson, R., Murphy, B. D., Renfree, M. B., et al. (2013). A new role for muscle segment homeobox genes in mammalian embryonic diapause. Open biology 3, 130035.

de Ziegler, D., Bulletti, C., Fanchin, R., Epiney, M. and Brioschi, P. A. (2001). Contractility of the nonpregnant uterus: the follicular phase. Annals of the New York Academy of Sciences 943, 172–184.

Douglas, D. A., Houde, A., Song, J. H., Farookhi, R., Concannon, P. W. and Murphy, B. D. (1998). Luteotropic hormone receptors in the ovary of the mink (Mustela vison) during delayed implantation and early-postimplantation gestation. Biol Reprod 59, 571–578.

Dupont, S., Krust, A., Gansmuller, A., Dierich, A., Chambon, P. and Mark, M. (2000). Effect of single and compound knockouts of estrogen receptors alpha (ERalpha) and beta (ERbeta) on mouse reproductive phenotypes. Development 127, 4277–4291.

Enders, A. C. and Buchanan, G. D. (1959). The reproductive tract of the female nine-banded armadillo. Texas reports on biology and medicine 17, 323–340.

Fenelon, J. C., Banerjee, A. and Murphy, B. D. (2014). Embryonic diapause: development on hold. Int J Dev Biol 58, 163–174.

Flores, D., Madhavan, M., Wright, S. and Arora, R. (2020). Mechanical and signaling mechanisms that guide pre-implantation embryo movement. Development 147.

Franco, H. L., Rubel, C. A., Large, M. J., Wetendorf, M., Fernandez-Valdivia, R., Jeong, J. W., Spencer, T. E., Behringer, R. R., Lydon, J. P. and Demayo, F. J. (2012). Epithelial progesterone receptor exhibits pleiotropic roles in uterine development and function. FASEB journal : official publication of the Federation of American Societies for Experimental Biology 26, 1218–1227.

Fu, Z., Wang, B., Wang, S., Wu, W., Wang, Q., Chen, Y., Kong, S., Lu, J., Tang, Z., Ran, H., et al. (2014). Integral proteomic analysis of blastocysts reveals key molecular machinery governing embryonic diapause and reactivation for implantation in mice. Biol Reprod 90, 52.

Ghosh, A., Syed, S. M., Kumar, M., Carpenter, T. J., Teixeira, J. M., Houairia, N., Negi, S. and Tanwar, P. S. (2020). In Vivo Cell Fate Tracing Provides No Evidence for Mesenchymal to Epithelial Transition in Adult Fallopian Tube and Uterus. Cell reports 31, 107631.

Gidley-Baird, A. A., O’Neill, C., Sinosich, M. J., Porter, R. N., Pike, I. L. and Saunders, D. M. (1986). Failure of implantation in human in vitro fertilization and embryo transfer patients: the effects of altered progesterone/estrogen ratios in humans and mice. Fertil Steril 45, 69–74.

Hamatani, T., Daikoku, T., Wang, H., Matsumoto, H., Carter, M. G., Ko, M. S. and Dey, S. K. (2004). Global gene expression analysis identifies molecular pathways distinguishing blastocyst dormancy and activation. Proc Natl Acad Sci U S A 101, 10326–10331.

He, B., Zhang, H., Wang, J., Liu, M., Sun, Y., Guo, C., Lu, J., Wang, H. and Kong, S. (2019). Blastocyst activation engenders transcriptome reprogram affecting X-chromosome reactivation and inflammatory trigger of implantation. Proc Natl Acad Sci U S A 116, 16621–16630.

Kastelic, J. P., Adams, G. P. and Ginther, O. J. (1987). Role of progesterone in mobility, fixation, orientation, and survival of the equine embryonic vesicle. Theriogenology 27, 655–663.

Kendle, K. E. and Lee, B. (1980). Investigation of the influence of progesterone on mouse embryo transport by using antiprogestational steroids. J Reprod Fertil 58, 253–258.

Laird, N. M. and Ware, J. H. (1982). Random-effects models for longitudinal data. Biometrics 38, 963–974.

Liang, Y. X., Liu, L., Jin, Z. Y., Liang, X. H., Fu, Y. S., Gu, X. W. and Yang, Z. M. (2018). The high concentration of progesterone is harmful for endometrial receptivity and decidualization. Sci Rep 8, 712.

Lopes, F. L., Desmarais, J. A. and Murphy, B. D. (2004). Embryonic diapause and its regulation. Reproduction 128, 669–678.

Lydon, J. P., DeMayo, F. J., Funk, C. R., Mani, S. K., Hughes, A. R., Montgomery, C. A., Jr., Shyamala, G., Conneely, O. M. and O’Malley, B. W. (1995). Mice lacking progesterone receptor exhibit pleiotropic reproductive abnormalities. Genes & development 9, 2266–2278.

Ma, W. G., Song, H., Das, S. K., Paria, B. C. and Dey, S. K. (2003). Estrogen is a critical determinant that specifies the duration of the window of uterine receptivity for implantation. Proc Natl Acad Sci U S A 100, 2963–2968.

Mackler, A. M., Green, L. M., McMillan, P. J. and Yellon, S. M. (2000). Distribution and activation of uterine mononuclear phagocytes in peripartum endometrium and myometrium of the mouse. Biol Reprod 62, 1193–1200.

Mantalenakis, S. J. and Ketchel, M. M. (1966). Frequency and extent of delayed implantation in lactating rats and mice. J Reprod Fertil 12, 391–394.

McCormack, J. T. and Greenwald, G. S. (1974). Evidence for a preimplantation rise in oestradiol–17beta levels on day 4 of pregnancy in the mouse. J Reprod Fertil 41, 297–301.

McLaren, A. (1968). A study of balstocysts during delay and subsequent implantation in lactating mice. J Endocrinol 42, 453–463.

McLaren, A. (1971). Blastocysts in the mouse uterus: the effect of ovariectomy, progesterone and oestrogen. J Endocrinol 50, 515–526.

Mead, R. A. (1981). Delayed implantation in mustelids, with special emphasis on the spotted skunk. Journal of reproduction and fertility. Supplement 29, 11–24.

Mead, R. A. (1993). Embryonic diapause in vertebrates. The Journal of experimental zoology 266, 629–641.

Nilsson, O. (1974). The morphology of blastocyst implantation. J Reprod Fertil 39, 187–194.

Norris, M. L. and Adams, C. E. (1981). Concurrent lactation and reproductive performance in CFLP mice mated post partum. Laboratory animals 15, 273–275.

Paria, B. C., Huet-Hudson, Y. M. and Dey, S. K. (1993). Blastocyst’s state of activity determines the “window” of implantation in the receptive mouse uterus. Proc Natl Acad Sci U S A 90, 10159–10162.

Pinehiro, J., Bates, D., DebRoy, S., Sarkar, D. and Team, R. C. (2021). nlme: Linear and Nonlinear Mixed Effects Models.

Psychoyos, A. (1965). [CONTROL OF NIDATION IN MAMMALS]. Archives d’anatomie microscopique et de morphologie experimentale 54, 85–104.

Psychoyos, A. (1973). Hormonal control of ovoimplantation. Vitamins and hormones 31, 201–256.

Renfree, M. B. and Shaw, G. (2014). Embryo-endometrial interactions during early development after embryonic diapause in the marsupial tammar wallaby. Int J Dev Biol 58, 175–181.

Restall, B. J. and Bindon, B. M. (1971). The timing and variation of pre-implantation events in the mouse. J Reprod Fertil 24, 423–426.

Roblero, L. S., Fernández, O. and Croxatto, H. B. (1987). The effect of RU486 on transport, development and implantation of mouse embryos. Contraception 36, 549–555.

Sebag-Peyrelevade, S. and Fanchin, R. (2015). Uterine Contractility and Embryo Transfer. In Human Embryo Transfer (ed. G. N. Allahbadia & C. F. Chillik).

Team, R. C. (2014). R: A Language and Environment for Statistical Computing.

Tourgeman, D. E., Boostanfar, R., Chang, L., Lu, J., Stanczyk, F. Z. and Paulson, R. J. (2001). Is there evidence for preferential delivery of ovarian estradiol to the endometrium? Fertil Steril 75, 1156–1158.

van Houten, E. L. and Visser, J. A. (2014). Mouse models to study polycystic ovary syndrome: a possible link between metabolism and ovarian function? Reprod Biol 14, 32–43.

Wang, H. and Dey, S. K. (2006). Roadmap to embryo implantation: clues from mouse models. Nat Rev Genet 7, 185–199.

Whitten, W. K. (1955). Endocrine studies on delayed implantation in lactating mice. J Endocrinol 13, 1–6.

Whitten, W. K. (1958). Endocrine studies on delayed implantation in lactating mice; role of the pituitary in implantation. J Endocrinol 16, 435–440.

Winuthayanon, W., Hewitt, S. C., Orvis, G. D., Behringer, R. R. and Korach, K. S. (2010). Uterine epithelial estrogen receptor alpha is dispensable for proliferation but essential for complete biological and biochemical responses. Proc Natl Acad Sci U S A 107, 19272–19277.

Yoshii, A., Kitahara, S., Ueta, H., Matsuno, K. and Ezaki, T. (2014). Role of uterine contraction in regeneration of the murine postpartum endometrium. Biol Reprod 91, 32.

Yoshinaga, K. (1961). Effect of local application of ovarian hormones on the delay in implantation in lactating rats. J Reprod Fertil 2, 35–41.

Yoshinaga, K. and Adams, C. E. (1966). Delayed implantation in the spayed, progesterone treated adult mouse. J Reprod Fertil 12, 593–595.

